# Increasing vitrification temperature improves the cryo-electron microscopy reconstruction

**DOI:** 10.1101/824698

**Authors:** Huigang Shi, Wuchun Ling, Dongjie Zhu, Xinzheng Zhang

## Abstract

At the initial stage of the cryo-electron microcopy (cryo-EM) samples irradiated by electrons, the cryo-EM samples suffer from a rapid “burst” phase (first 3~4 e^−^/Å^2^) of beam induced motion (BIM) which is too fast to be corrected by existing motion correction software, and lowers the quality of the initial frames. Therefore, these least radiation damaged, but ruined frames are commonly excluded or down-weighted during data processing, which reduces the undamaged signals in the reconstruction and decreases the reconstruction resolution by weakening the averaging power. Here, we show that increasing the freezing temperature of cryo-EM samples suppresses the BIM in this phase. The quality of initial frames is partially recovered after BIM correction and is better than that of subsequent frames in certain frames. Incorporating these initial frames into the reconstruction increases the resolution, at an equivalent of ~60% extra data. Moreover, these frames are least radiation damaged, thus preserves the high quality cryo-EM density of radiation sensitive residues. Such density is usually damaged or very weak in the canonical 3D reconstruction. In addition, we found that a different kind of radiation damage neglected previously occurs in the per-frame reconstruction after the exposure of 2.5 e^−^/Å^2^. Such radiation damage distorts the density of atoms. The deformation can be avoided by only including the frames from the first 2.5 e^−^/Å^2^ into the reconstruction. Overall, the high temperature freezing not only provides extra undamaged signal to the reconstruction, but also increases the resolution of the reconstruction.

## Introductions

A protein sample is structurally damaged when it is irradiated by electrons during electron microscopy. The high energy electrons first break chemical bonds in either proteins or the surrounding buffers, which produces free radicals. These radicals then cause a cascade of chemical reactions to further damage the protein structure. The freezing of the protein sample can help to reduce such radiation damage ^1,2^. At liquid nitrogen temperature, the free radicals are “trapped” locally and their mobility is greatly reduced, which results in less chemical reactions and limits the radiation damage ^3^. Currently, the protein sample is embedded in vitreous ice through a cryo-EM sample preparation procedure and then imaged at ~−180 °C to reduce the radiation damage.

It has been found that the vitreous ice could form by plunge freezing a microdroplet at ~−160 °C into liquid ethane and turned to multicrystal when its temperature was raised to ~−135 °C ^4^. Currently, a similar cryo-EM sample preparation procedure is performed using either home-made plunge devices or other automatic freezing devices by plunging the cryo-EM grid into liquid ethane maintained at ~−180 °C to avoid crystal ice ^5^.

During the imaging of the vitrified cryo-EM sample, the electron irradiation causes BIM of the proteins. BIM is usually characterized by two phases, the initial rapid “burst” phase and a subsequent slower phase ^5–7^. The cause of the rapid “burst” phase is unknown. Possible explanations for the rapid “burst” phase are that at the beginning of the electron irradiation, the stress in the vitreous ice preserved during fast freezing or generated by initial radiation damage is released ^8^, which causes the motion. The slower phase of BIM may be caused by the radiation damage of the vitreous ice or protein sample and also by the beam-induced Brownian motion of water molecules ^9^. The BIM blurs the cryo-EM images and results in a decrease in resolution of the reconstruction. Using the movie mode of the direct electron detector for imaging, a dominating part of BIM in the slower phase can be corrected by translational aligning the frames of a movie ^10–13^. However, the BIM in the rapid “burst” phase is probably too fast to be effectively corrected. Thus, the resolution of the per-frame reconstructions calculated by the frames exposed to the first 3 or 4 e^−^/Å^2^ within the rapid “burst” phase is lower than that of the per-frame reconstructions reconstructed by the subsequent frames. Currently, frames suffered from rapid “burst” phase are either excluded or down-weighted in further data process. When cryo-EM samples are exposed to electrons, the structure of the protein is irreversibly and gradually damaged by the accumulating electrons. Less and less structural information is preserved in the succeed frames. For instance, only about one third of 3 Å spatial frequency signal is preserved in the sample after the exposure of 4 e^−^/Å ^2^ ^14^. Therefore, the discarded or down-weighted frames containing the most high frequency signals are essential for high resolution reconstruction.

Here, we show that different protein samples, such as apo-ferritin, GDH, hemoglobin and so on can be vitrified by plunge freezing the cryo-EM grid into liquid ethane at −110°C. The rapid “burst” phase of BIM weakened along with the increasing of freezing temperature. The initial frames can be recovered from BIM and incorporated into the data to improve the resolution and enhance the undamaged signals in the cryo-EM map.

## Results and discussions

### Vitrification of cryo-EM sample at different temperatures

Fast freezing may cause the stress preserved in the cryo-EM sample. The vitrification at high temperature may lead to high mobility of the atoms during vitrification and reduce the stress preserved in the vitreous ice. To test the ability of vitrifying cryo-EM sample at different temperatures, a thin layer of aqueous sample containing apo-ferritin was frozen by plunging the grid into liquid ethane at different temperatures ranging from −90 °C to −180 °C using automatic plunging devices (Gatan CP3 and Leica EMGP). To accurately monitor the temperature of liquid ethane in automatic plunging devices, we used a temperature-measuring thermocouple to measure the temperature of liquid ethane in both CP3 and EMGP (See SI Appendix, Table. S1 and methods for details). We further calibrated the temperature by measuring the liquid nitrogen and the melting points of solid ethane, solid propane and ice at atmospheric pressure and comparing them with the previously reported melting points as shown in Supplementary Fig. 1 and Supplementary Tab. 1. Based on the calibrated temperature, our freezing experiments showed that the thin layer of aqueous sample can be vitrified at a temperature below −110 °C in liquid ethane. At a temperature above −110 °C, the crystal ice dominated the frozen sample. As shown in Supplementary Fig. 2, the crystal ice produced polycrystalline diffraction which can be easily differentiated from the diffraction pattern of the vitreous ice. Such result was different from previously observations indicating that the microdroplet turned to multi-crystal when its temperature was raised to ~−135°C. The aqueous layer in our experiment has a thickness of less than 50 nm and may have a much faster cooling rate than that of a microdroplet frozen in liquid ethane. The difference in the cooling rate may affect the ability of vitrification ^15^. The apo-ferritin samples frozen at −180 °C, −145 °C and −110 °C in liquid ethane were stored in liquid nitrogen for further cryo-EM data collection.

### High vitrification temperature improves the resolution of the per-frame reconstructions

We collected a dataset on each sample mentioned above using a Titan Krios microscope equipped with K2 camera, where the sample was maintained at ~−193 °C. The apo-ferritin in the three datasets was reconstructed to 2.20 Å, 2.38 Å and 1.89 Å, respectively, employing a conventional data processing procedure of single particle analysis (details shown in methods). In addition, apo-ferritin particles from each frame were extracted from the aligned movie stacks and the per-frame reconstruction was calculated according to the alignment in the last iteration of the refinement. The resolutions of the per-frame reconstructions from three datasets were calculated as shown in Fig. 1. The frame was exposed to 1.2 e^−^/ Å^2^, 1.2 e^−^/ Å^2^ and 1.27 e^−^/ Å^2^, respectively. In the dataset of apo-ferritin frozen at −180 °C, the resolution of the first three per-frame reconstructions are significantly lower than that of the fourth per-frame reconstruction. Such difference becomes smaller with the increasing of the freezing temperature in the other two datasets. In the sample frozen at −110 °C, the per-frame reconstruction of the second frame achieved the best resolution, while the resolution of the per-frame reconstruction of the first frame was slightly lower.

**Fig. 1.**
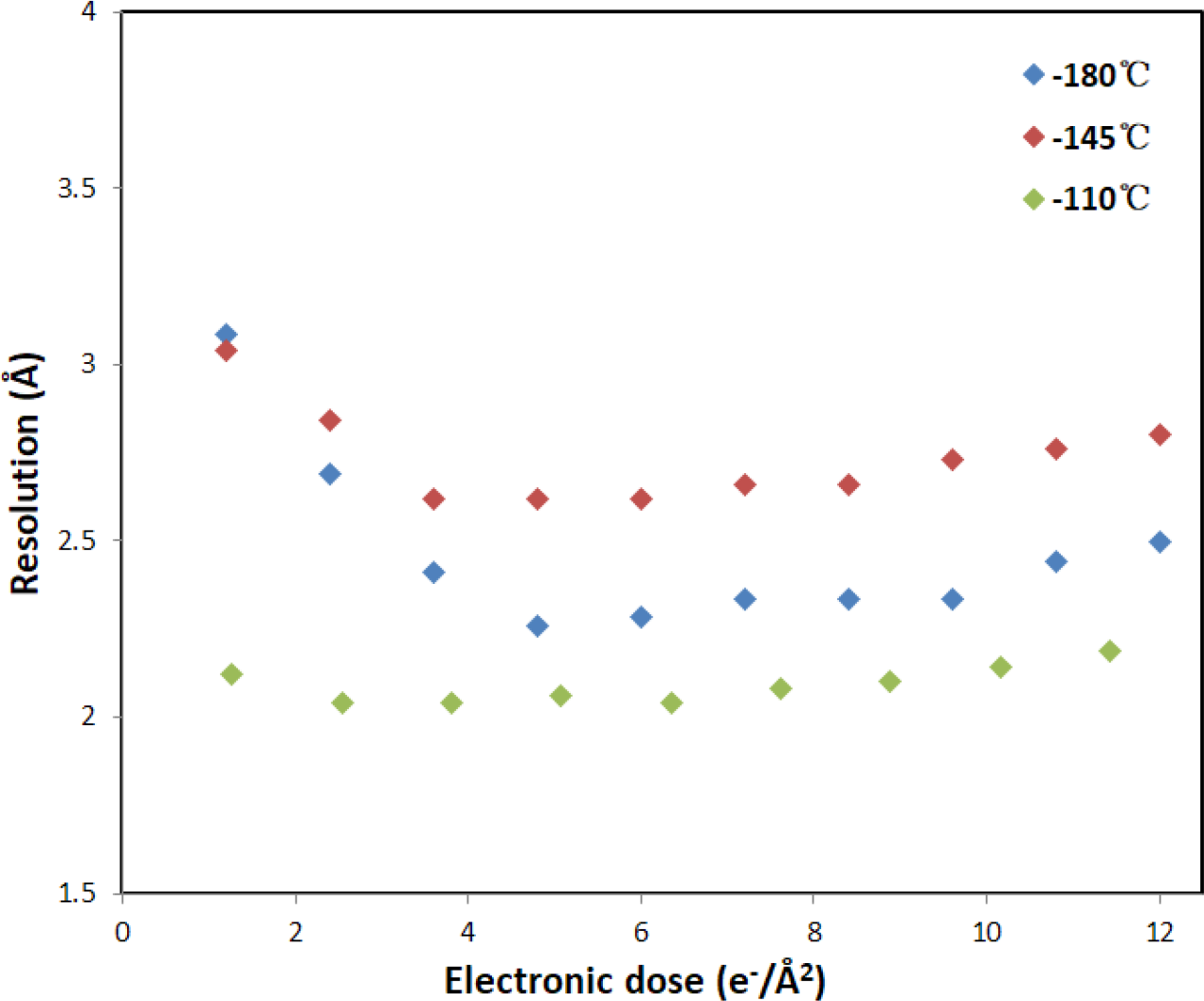
Resolution of per-frame reconstructions depending on the freezing temperatures. For the sample frozen at −180 °C, the regular temperature for cryo-EM sample preparation, the resolution of the per-frame reconstruction of the first four frames was decreased in an order (blue plot). The differences in resolution between the first three frames and the fourth frame became smaller when the freezing temperature increased to −145 °C (red plot). At the freezing temperature of −110 °C, the resolutions of the first three frames were similar to that of the following frame, where the second frame produced a per-frame reconstruction with the best resolution among all the frames (green plot).

Our results show that the qualities of the initial few frames become better with the increasing of freezing temperatures, which indicates that the high freezing temperature depresses the rapid “burst” phase of BIM. Therefore, it is very possible that the rapid “burst” phase is associated with the releasing of the stress frozen into the specimen during the plunge-freezing.

### The radiation damages in frames with different accumulated electrons

Certain residues in protein that contain side-chain carboxyl groups, solvent-exposed disulfide bonds or the active site of an enzyme are especially sensitive to radiation damage ^8,16^. We compared cryo-EM densities of site-specific amino acids in the different per-frame reconstructions. As shown in Fig. 2a, the side chain densities of radiation sensitive amino acids preserved well in the second per-frame reconstruction while these densities were damaged in the fourth or the fifth frame. The corresponding accumulated electron dosages before taking the fourth and fifth frames are 3.81 e^−^/Å^2^ and 5.08 e^−^/Å^2^, respectively. Since the conventional cryo-EM sample frozen at −180 °C only produces good per-frame reconstructions after the first 3 or 4 electrons, the dose-weighting procedure of the conventional cryo-EM sample preparation led to damaged side chain densities of these amino acids in the cryo-EM map as shown in Fig. 2a. The cryo-EM sample frozen at −110 °C produced much better densities of these radiation sensitive amino acids by averaging the first three undamaged frames.

**Fig. 2.**
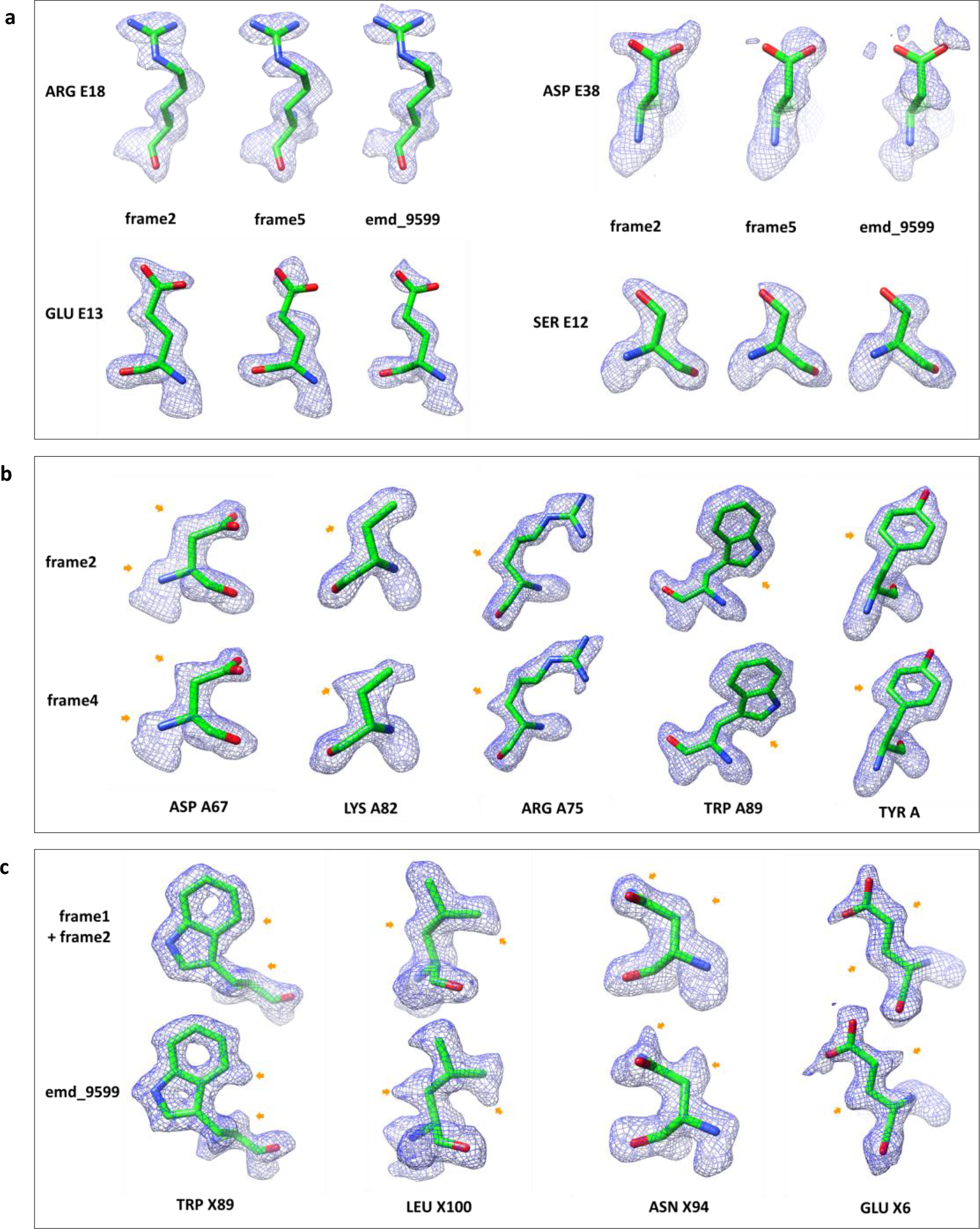
Radiation damages shown in per-frame reconstructions with different accumulated electron dosages. **a,** Electron densities from the radiation sensitive amino acids. The electron densities of amino acids in the second per-frame reconstruction showed an intact structure of the amino acid (left), while the side chain densities were damaged in the fifth per-frame reconstruction with a accumulated electron dosage of ~6.35 e^−^/Å^2^ (middle). The corresponding amino acids in the reconstruction (EMD_9599) of apo-ferritin frozen at ~−180°C showed weak or ruined side chain densities (right). **b,** In the fourth per-frame reconstruction, the electron densities of some of the atoms were deformed by extended to one direction as pointed by orange arrows. The corresponding amino acids in the second per-frame reconstruction showed in the upper row exhibited normal appearance. **c,** The electron densities of previously published apo-ferritin by conventional freezing with freezing temperature of ~−180 °C exhibited the same deformation. The deformed atoms from EMD _9599 were indicated by arrows as shown in the lower row. The electron densities of corresponding atoms in our reconstruction composed of the first two frames were shown in the upper raw.

In addition to the traditional radiation damage reducing the high resolution signals, we found a new type of radiation damage that deforms the cryo-EM density of atoms. As shown in Fig. 2b, compared with the cryo-EM densities in the per-frame reconstruction by the second frame, the cryo-EM densities of some atoms in the per-frame reconstructions by the third and fourth frames (in the range from ~2.5 to 5 e^−^/Å^2^) were extended in one direction. Such deformation disappeared in the succeeding frames, where the traditional radiation damage dominated and damaged the high resolution features. Such deformation occurred to quite a few atoms, and behaved similar in the two half-dataset maps for the calculation of Fourier Shell Correlations (FSC), therefore it was not detected by FSC and did not change the signal to noise ratio (SNR) of the map. The conventional SNR based dose-weighting procedure cannot reduce the deformation in the map. Only the reconstruction made by the first 2 frames can overcome the deformation problem (Fig. 2c), bearing a slightly decrease of the overall resolution.

Frames exposed to the first 3 or 4 e^−^/Å^2^ from cryo-EM sample frozen at −180 °C suffer from uncorrected BIM and are usually excluded or down-weighted from the data processing. The subsequent frames contribute the most of the high frequency signals to the reconstruction. As a result, the high resolution structural features of a high resolution reconstruction retain the deformation. Such deformation can be seen in different published high resolution maps calculated by single particle analysis as shown in Fig. 2c.

The cause of this new type of radiation damage is unknown. A possible explanation is that in the dose range from 2.5 to 5 e^−^/Å^2^, the incident electrons knock out a few electrons surrounding the atomic nucleus, which may cause positive charge on atoms. The resulting net charge may then cause the corresponding atom being away from the original position.

### High vitrification temperature improves the overall resolution

Using all frames processed by dose-weighting procedure in motionCorr2 (all-frames dataset), our apo-ferritin sample frozen at −110 °C led to an overall resolution of 1.89 Å. To compare the quality of images from our data (full frames dataset) and from the conventional cryo-EM data suffering from the rapid burst phase of BIM, we mimicked conventional cryo-EM data by excluding the first 3 frames from our data during the dose-weighting procedure. Using the same selected apo-ferritin particles, the resulting new dataset (partial-frames dataset) achieved an overall resolution of 1.97 Å. The *ResLog* plots from these two datasets indicate that in the partial-frames datasets mimicking the cryo-EM sample frozen at −180 °C, 60% additional particles are needed to achieve the same resolution as in the all-frames dataset corresponding to cryo-EM sample frozen at −110 °C (Fig. 3). Moreover, we observed a slight improvement of the b-factor in the all-frames dataset. The all-frames dataset exhibited higher image quality compared with the partial-frames dataset as the former has been averaged with the extra 3 frames with the least radiation damage. The higher image quality resulted in a more accurate alignment in the refinement during data processing, thus improving the b-factor.

**Fig. 3.**
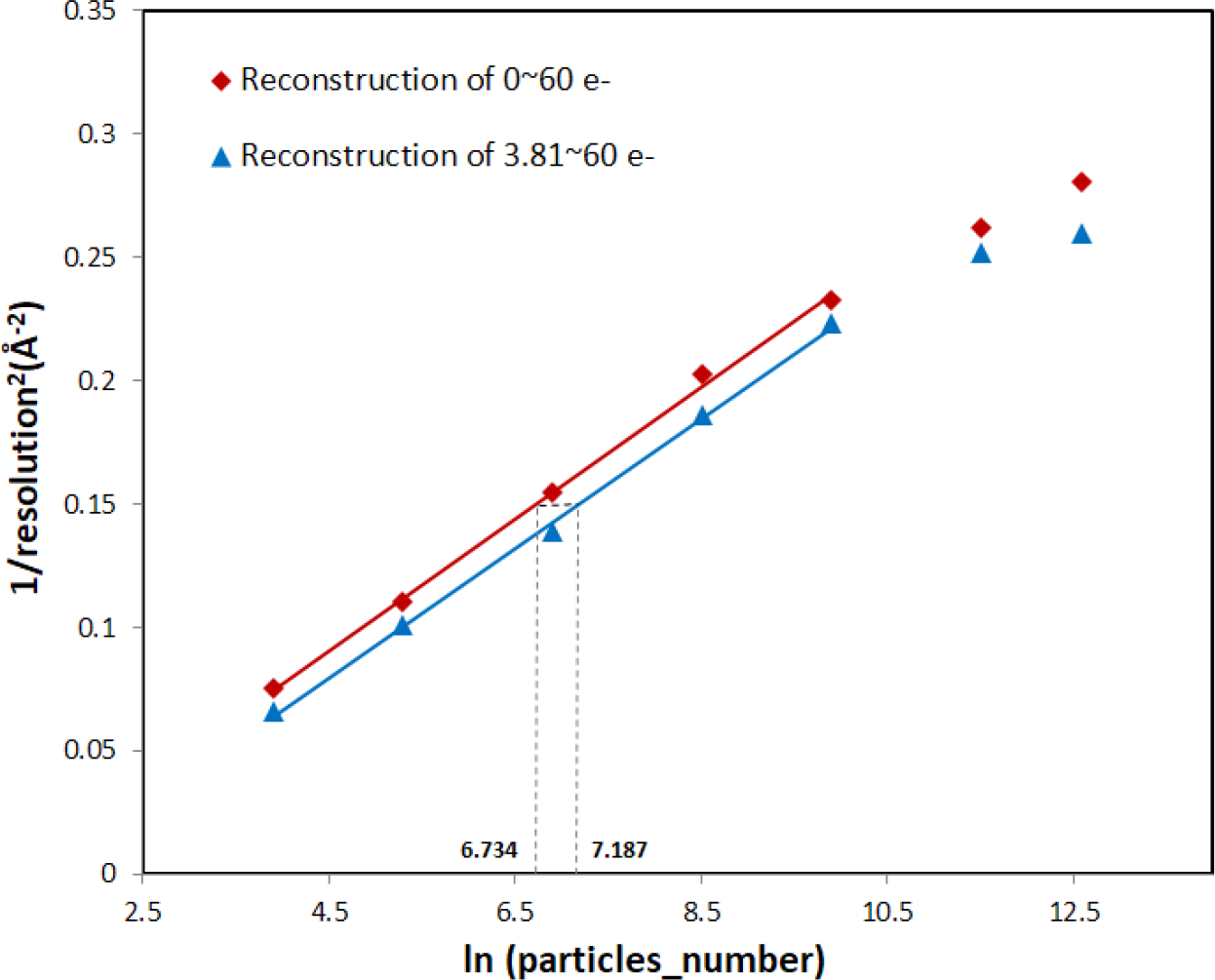
The *ResLog* plots of dataset including or excluding the first 3 frames. The red and blue straight lines were fitted from the linear part of the plot, corresponding to the dataset with or without the first 3 frames. Using the same number of apo-ferritin particles, the dataset with the first 3 frames achieved a higher resolution than the dataset without the first 3 frames. The horizontal dash line in the middle of the linear region is 0.453 in length, which indicated that in the dataset without the first 3 frames, ~1.6× particles had to be included in the reconstruction to achieve the same resolution as in the dataset with the first 3 frames.

### Vitrification at high temperature can be widely applied to different cryo-EM samples

We have tested the freezing on other protein samples, such as GDH (Supplementary Fig. 3), hemoglobin (Supplementary Fig. 5a) and Cas3-AcrF3 (130 kDa protein, Supplementary Fig. 5b), that can be frozen into a thin layer of ice. The method of plunge freezing at −110 °C produced identical vitreous ice around these samples, suggesting that this method can be applied to different proteins. We collected cryo-EM data and performed per-frame reconstruction on the GDH sample frozen at −110 °C, which behaved similar to the apo-ferritin sample frozen at −110 °C (Supplementary Fig. 3).

We also tested to freeze a virus sample with large diameter of ~35 nm at −110 °C. In such a case, a thicker layer of ice (~50 nm) is required for embedding the virus. The freezing of the virus at −110 °C led to crystal ice probably due to the decrease of the cooling rate caused by the increasing of the thickness of the aqueous sample. When the freezing temperature was further decreased to −135 °C, we were able to vitrify this virus sample and other virus sample with larger diameters (Supplementary Fig. 5c and Supplementary Fig. 5d). The cryo-EM data was collected on this sample and per-frame reconstructions were calculated. The resolutions of per frame reconstructions were compared with that of the virus frozen at −180 °C. As shown in Supplementary Fig. 4, the difference in resolutions between the per-frame reconstructions by initial frames and the best per-frame reconstruction became smaller in the virus frozen at −135 °C compared with that of the virus frozen at −180 °C. It indicates that the increasing of the freezing temperature of a virus sample can only partially recover resolutions of reconstructions by initial frames. However, the partial recovery of the initial frames should still benefit the final reconstruction.

Based on the result above, it seems that the rapid burst phase of BIM is associated with the freezing temperature more than with the cooling rate. Therefore, the finding of a better coolant or different techniques of cooling that increases the cooling rate of the protein sample would help to increase the temperature of vitrification and completely eliminate the rapid burst phase of BIM in cryo-EM.

## Conclusions

We demonstrated that a thin layer of protein sample can be vitrified at a temperature below −110 °C. By analyzing the cryo-EM data of protein samples vitrified at different temperatures, we showed that the increasing of freezing temperature suppresses the rapid burst phase of BIM. Thus, the quality of frames at this stage can be improved efficiently from the BIM, which leads to an increasing of the resolution of the per-frame reconstructions. The resolution of the overall reconstruction is improved by Incorporating the high quality initial frames, which is equivilent to the adding of extra 60% data. The improvement of the per-frame reconstruction by initial frames allows the reconstruction of densities of intact radiation sensitive amino acids before radiation damage. In addition, we identified a new type of radiation damage, which only occurs at the early stage of electron irradiation. Such radiation damage deforms the densities of amino acids and can only be avoided by including the frames exposed by first 2.5 e^−^/Å^2^ into the reconstruction. Overall, the increasing of the freezing temperature in cryo-EM specimen preparation can not only improve the resolution of reconstruction, but also provide extra undamaged signal to the reconstruction.

## Methods

### Protein and cryo-EM specimen preparation

The human apo-ferritin ^17,18^ (a gift from Prof. Yan’s lab) was diluted to a concentration of ~2 mg/ml. The GDH (Sigma-Aldrich, catalog #G2626) was dialyzed in 100 mM potassium phosphate [pH 6.8] overnight prior to purification by Gel filtration ^19^. The concentration of GDH for cryo-EM sample preparation was ~3 mg/ml.

Approximately 2 μl aliquots of apo-ferritin sample were applied to each glow-discharged NiTi foil R1.2/1.3(Au) grid. Grids were flash plunged into liquid ethane at different temperatures ranging from −90 °C to −180 °C after being blotted for 5s by automatic plunging device CP3.

Approximately 3 μl aliquots of GDH, hemoglobin and a Cas3-AcrF3 (130 kDa protein) were applied to glow-discharged GIG R2/1(Cu) grids. These grids were flash plunged into liquid ethane at different temperatures after being blotted for 5s by automatic plunging device CP3.

Approximately 3 μl aliquots of virus with diameter of ~35 nm, ~55 nm and ~95 nm (a gift from Prof. Gao’s lab) were applied to glow-discharged NiTi foil R1.2/1.3 (Au) grids. These grids were flash plunged into liquid ethane at different temperatures after being blotted for 4s by automatic plunging device EMGP.

### Data acquisition and processing

Three datasets of apo-ferritin frozen at −180 °C, −145 °C and −110 °C were collected in FEI Titan Krios G2 equipped with direct detector K2 summit (super resolution mode) and GIF quantum energy filter (energy width of 20 eV) at magnifications of 165,000, 165,000 and 265,000, yielding binned pixel sizes of 0.827 Å, 0.827 Å and 0.515 Å, respectively. The corresponding dose rates for three datasets were measured as ~12 e^−^/Å^2^/s, 12 e^−^/Å^2^/s and 25.1 e^−^/Å^2^/s and the exposure times were 5 s, 5 s and 2.39 s, respectively. The images were fractionated into 50, 50 and 47 frames, respectively. 567, 542 and 1,542 micrographs were collected for three datasets, respectively. The defocus of collected micrographs ranged from 0.5 μm to 3 μm. Two datasets of GDH samples frozen at −180 °C and −110 °C were collected in FEI Titan Krios G2 equipped with direct detector K2 summit (super resolution mode) and GIF quantum energy filter (energy width of 20 eV) at a magnification of 165,000, yielding a binned pixel size of 0.827 Å, with a dose rate of ~12.1 e^−^/Å^2^/s and the exposure time of 4.13 s. The micrographs were fractionated into 50 frames. 1154 and 1274 micrographs were collected for two datasets, respectively. The defocus ranged from 0.5 μm to 3 μm. The two datasets of 35 nm virus samples frozen at −180 °C and −135 °C was collected in FEI Titan Krios G2 equipped with direct detector K2 summit (super resolution mode) at a magnification of 130,000 and 165,000, with a dose rate of ~9.24 e^−^/Å^2^/s and 13.5 e^−^/Å^2^/s and the exposure time of 6.5 s and 3.7 s, yielding a binned pixel size of 1.050 and 0.827 Å, respectively. 602 and 369 micrographs were collected for two datasets, respectively. The defocus range was from 0.5 μm to 3 μm.

In all datasets, the BIM was corrected by MotionCor2 ^10^. The aligned stacks were saved for subsequent per-frame reconstructions. The defocus parameters were estimated by CTFFIND4 ^20^. About 10000 particles were picked by e2boxer.py (a semiautomatic particle picking software) ^21^ and were processed by 2D classification, yielding featured 2D averages. These featured 2D averages were used as references for automatic particle picking from all micrographs using RELION ^22^. After reference-free 2D classification and 3D classification, 164,335, 76,574 and 296,695 particles were kept for further refinement, yielding three reconstructions with resolutions of 2.37 Å, 2.56 Å and 2.03 Å, respectively. After performing the CTF refinement in RELION-3 ^23^, the resolution of reconstructions reached 2.20, 2.38 and 1.89 Å, respectively. The resolution of reconstruction was determined by post process in RELION. The same mask was used in the post process for all reconstructions of apo-ferritin. For GDH samples frozen at −180 °C and −115 °C, 83,207 and 179,441 particles from 1154 and 1274 micrographs were kept after 2D and 3D classification for 3D refinement, yielding 2.84 Å and 2.57 Å reconstructions, respectively. For 35 nm virus samples frozen at −180 °C and −135 °C, 18,881 and 3,419 particles from 602 and 369 micrographs were selected after 2D and 3D classifications, respectively. The further refinements yielded 2.97 Å and 3.25 Å reconstructions, respectively.

The single frame was split from the aligned stacks. Per-frame reconstructions were calculated from frames with parameters from the last iteration of refinement by *relion_reconstruct_mpi*. The resolution of per-frame reconstruction was calculated by post process in RELION applying the same soft mask for each sample.

### Atomic Model Refinement

The PDB coordinates 1MFR was served as a starting model for building the atomic model of apo-ferritin. Amino acids were mutated in COOT ^24^ according to the amino acids sequence of our sample. Manual adjustments were made in COOT. The model was refined by Phenix ^25^.

### The calibration of freezing temperatures in EMGP and CP3

We used a temperature thermocouple to calibrate the freezing temperatures reported in EMGP and CP3. First, we calibrated the temperature thermocouple by measuring the melting points of water, ethane and propane by putting the temperature thermocouple in the corresponding solid-liquid states. The deviation of output temperatures from melting points previously reported was calibrated by an exponential function (Supplementary Fig. 1). After the compensation of the deviation, the temperature thermocouple was used to measure the temperature of the liquid ethane in EMGP and CP3 at the location where the sample is frozen. The measured temperatures by the thermocouple after the compensation were compared to the reported temperatures in both EMGP and CP3 as shown in SI Appendix, Table. S1. The reported temperatures of EMGP are closer to the measured temperatures than that of CP3. We found the temperature of liquid ethane increased from the top to the bottom of the liquid ethane-containing cup in CP3, which indicates that the heating component controlling the temperature is located near the bottom of the cup. It is possible that the temperature sensor of CP3 is placed at the bottom of the cup, which may partially result in such a difference between measured temperature and reported temperature. Thus, we used the reported temperatures from EMGP as a standard in the whole study.

## Acknowledgement

We thank Prof. K.L. Fan and Prof. X.Y. Yan at the Institute of Biophysics, Chinese Academy Sciences, Beijing, China for preparing the apo-ferritin sample. We thank Y.Z. Cui, Dr. N. Du and Prof. GF. Gao at Institute of Microbiology, Chinese Academy of Sciences, Beijing, China for preparing virus sample. We thank Dr. J. Ma and Prof. Y.L. Wang at the Institute of Biophysics, Chinese Academy Sciences, Beijing, China for preparing Cas3-AcrF3 sample. We thank L. Kong for cryo-EM data storage and backup. Cryo-EM data collection was carried out at the Center for Biological Imaging, Core Facilities for Protein Science at the Institute of Biophysics (IBP), Chinese Academy of Sciences (CAS). We thank G. Ji, B.L. Zhu, F. Sun, and other staff members at the Center for Biological Imaging (IBP, CAS). We thank P.Y. Xia and Y.Z. Ma from the State Key Laboratory of Membrane Biology, Institute of Zoology, Chinese Academy of Sciences, for their support with EM. The project was funded by the National Key R&D Program of China (2017YFA0504700) to X.Z., and X.Z. received scholarships from the National Thousand (Young) Talents Program from the Office of Global Experts Recruitment in China.

## Author Contributions

X.Z. designed the experiment. H.S. and C.W. performed the experiments. H.S. and C.W. prepared the cryo-EM specimen and performed data collection. H.S., C.W. and D.Z. performed data processing. H.S., C.W. and X.Z. analyzed the result and wrote the manuscript. All authors discussed and commented on the results and the manuscript.

## Supplementary Figures

**Fig. S1.**
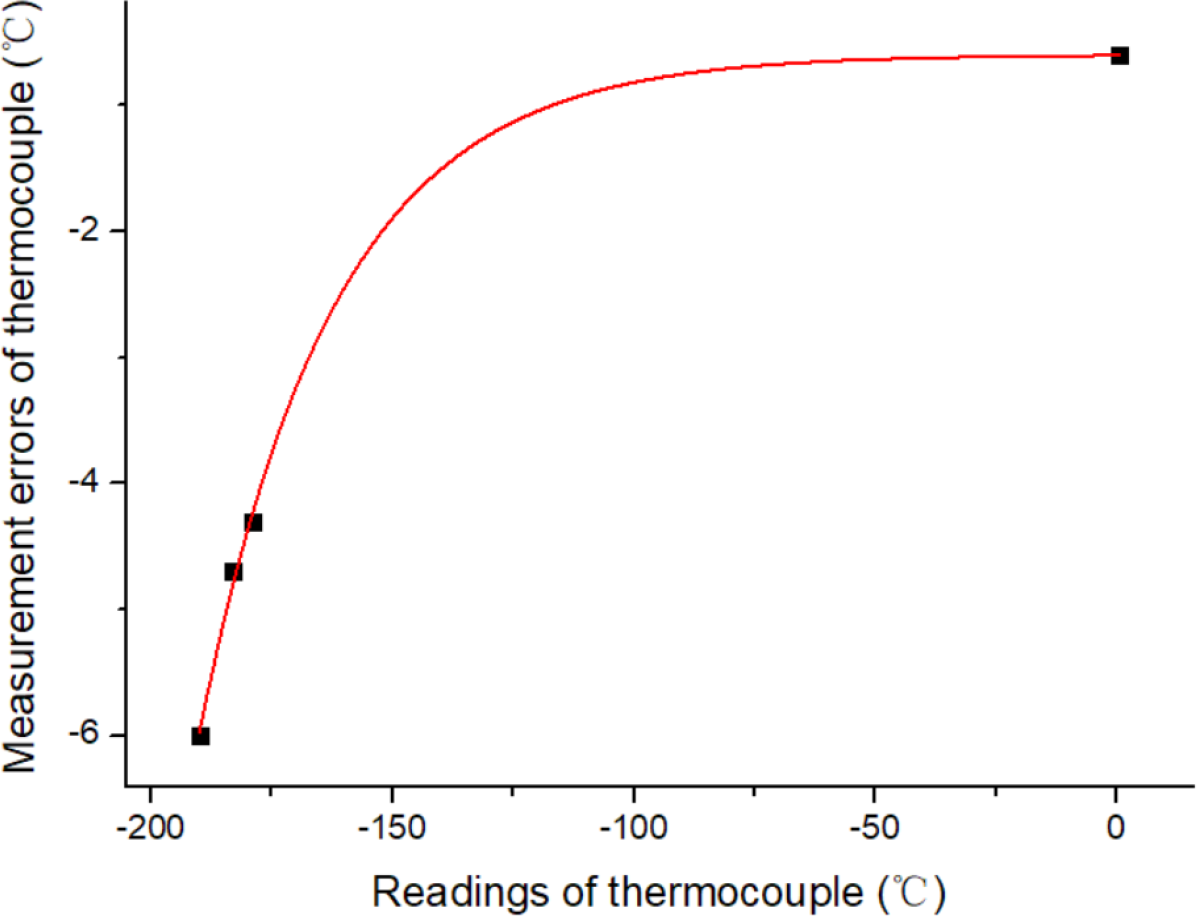
The plot of temperature thermocouple measurement errors. The differences in temperatures between measured temperatures of the solid-liquid phase of water, propane and ethane and the liquid nitrogen and the corresponding previously reported temperatures (0 °C for solid-liquid phase of water; −187.7 °C for solid-liquid phase of propane; −183.3 °C for solid-liquid phase of ethane and −196 °C for liquid nitrogen) at an atmospheric pressure, respectively. Four data points were well fitted to an exponential function (red).

**Table. S1.**
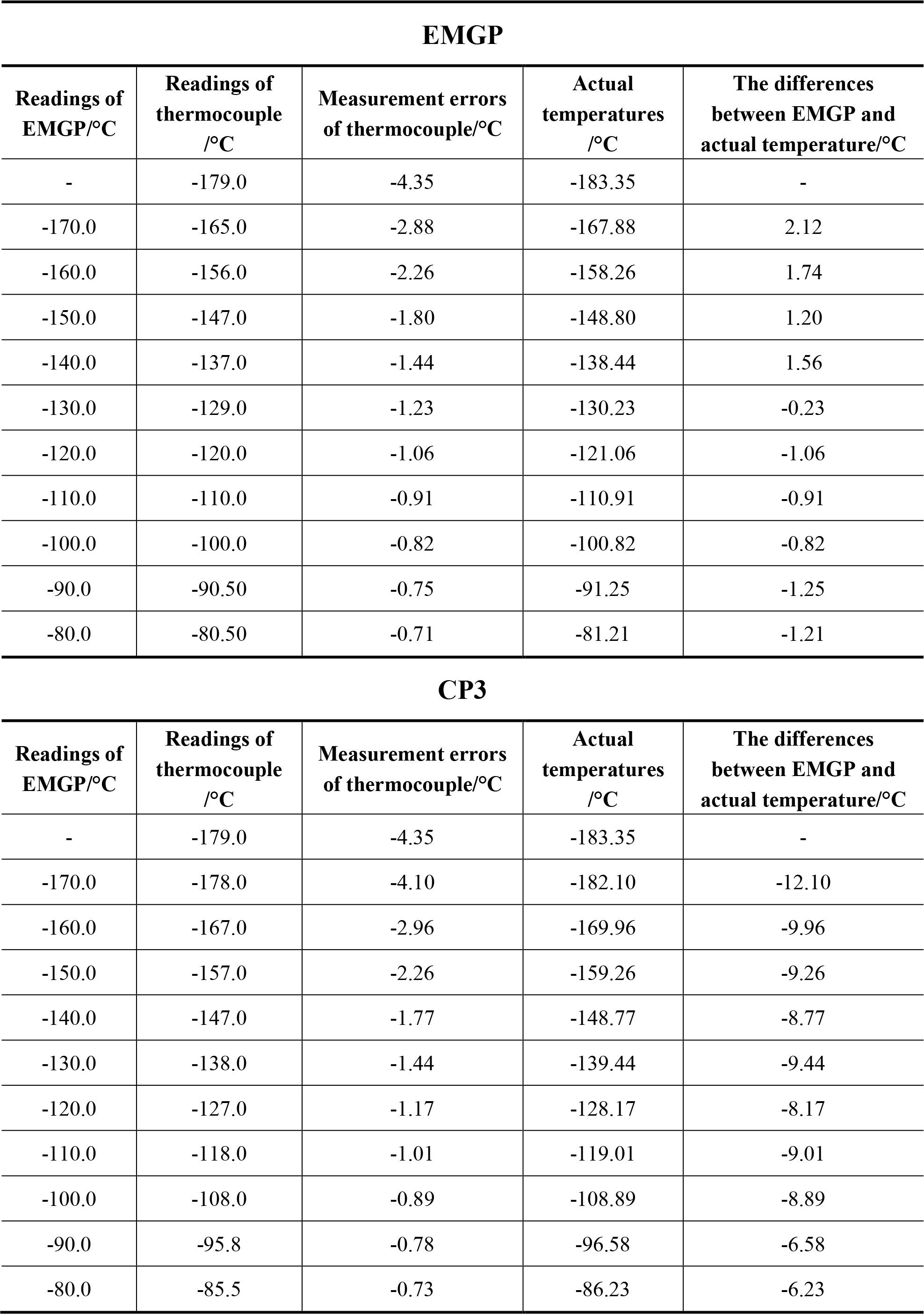
The temperature calibration of EMGP and CP3 using the temperature thermocouple.

**EMGP**
| Readings of EMGP/ °C | Readings of thermocouple / °C | Measurement errors of thermocouple/ °C | Actual temperatures / °C | The differences between EMGP and actual temperature/ °C |
| --- | --- | --- | --- | --- |
| - | -179.0 | -4.35 | -183.35 | - |
| -170.0 | -165.0 | -2.88 | -167.88 | 2.12 |
| -160.0 | -156.0 | -2.26 | -158.26 | 1.74 |
| -150.0 | -147.0 | -1.80 | -148.80 | 1.20 |
| -140.0 | -137.0 | -1.44 | -138.44 | 1.56 |
| -130.0 | -129.0 | -1.23 | -130.23 | -0.23 |
| -120.0 | -120.0 | -1.06 | -121.06 | -1.06 |
| -110.0 | -110.0 | -0.91 | -110.91 | -0.91 |
| -100.0 | -100.0 | -0.82 | -100.82 | -0.82 |
| -90.0 | -90.50 | -0.75 | -91.25 | -1.25 |
| -80.0 | -80.50 | -0.71 | -81.21 | -1.21 |

**CP3**
| Readings of EMGP/ °C | Readings of thermocouple / °C | Measurement errors of thermocouple/ °C | Actual temperatures / °C | The differences between EMGP and actual temperature/ °C |
| --- | --- | --- | --- | --- |
| - | -179.0 | -4.35 | -183.35 | - |
| -170.0 | -178.0 | -4.10 | -182.10 | -12.10 |
| -160.0 | -167.0 | -2.96 | -169.96 | -9.96 |
| -150.0 | -157.0 | -2.26 | -159.26 | -9.26 |
| -140.0 | -147.0 | -1.77 | -148.77 | -8.77 |
| -130.0 | -138.0 | -1.44 | -139.44 | -9.44 |
| -120.0 | -127.0 | -1.17 | -128.17 | -8.17 |
| -110.0 | -118.0 | -1.01 | -119.01 | -9.01 |
| -100.0 | -108.0 | -0.89 | -108.89 | -8.89 |
| -90.0 | -95.8 | -0.78 | -96.58 | -6.58 |
| -80.0 | -85.5 | -0.73 | -86.23 | -6.23 |

**Fig. S2.**
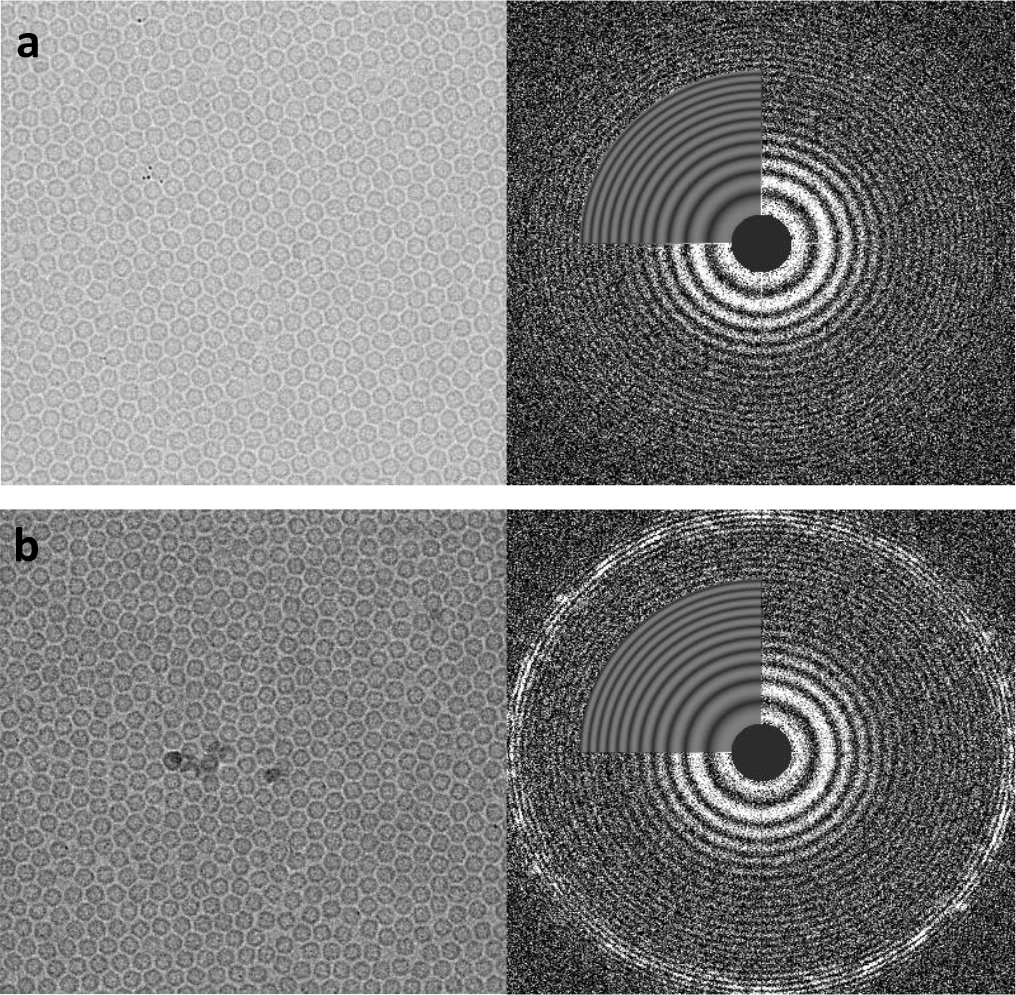
Cryo-EM Specimen embedded in vitreous ice and crystal ice. **a**, Apo-ferritin frozen at −110 °C was embedded in vitreous ice (left) and the corresponding Fourier transform (right). **b**, Apo-ferritin frozen at −90 °C was embedded in crystal ice (left) and the corresponding Fourier transform showing strong polycrystalline diffraction rings of ice at approx. 1/3.6 Å^−1^.

**Fig. S3.**
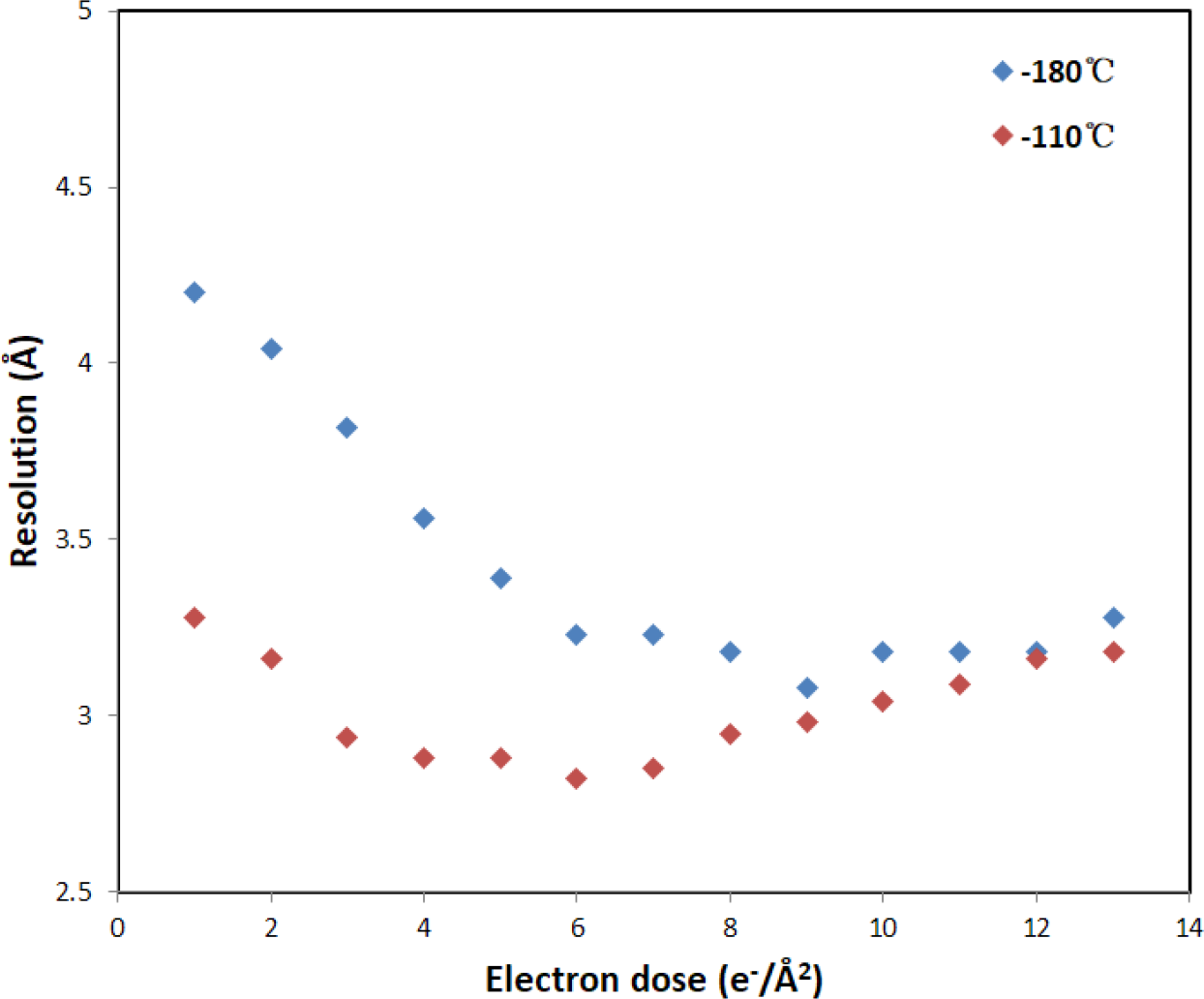
Resolution of per-frame reconstructions of GDH frozen at different temperatures. Each frame was exposed by ~1 e^−^/Å^2^ in both datasets. When the sample was frozen at −180 °C, the resolution of the per-frame reconstruction of the first five frames decreased significantly in reversed order. The following 7 frames produced reconstructions with similar resolutions (blue plot). While the sample was frozen at −110 °C, only the resolution of first two frames decreased significantly. The following 5 frames produced reconstructions with similar resolutions (red plot). The resolution of the reconstruction of the first frame is ~1.1 Å lower than that of the best reconstruction for the sample frozen at −180 °C while it is ~0.4 Å for the sample frozen at −110 °C.

**Fig. S4.**
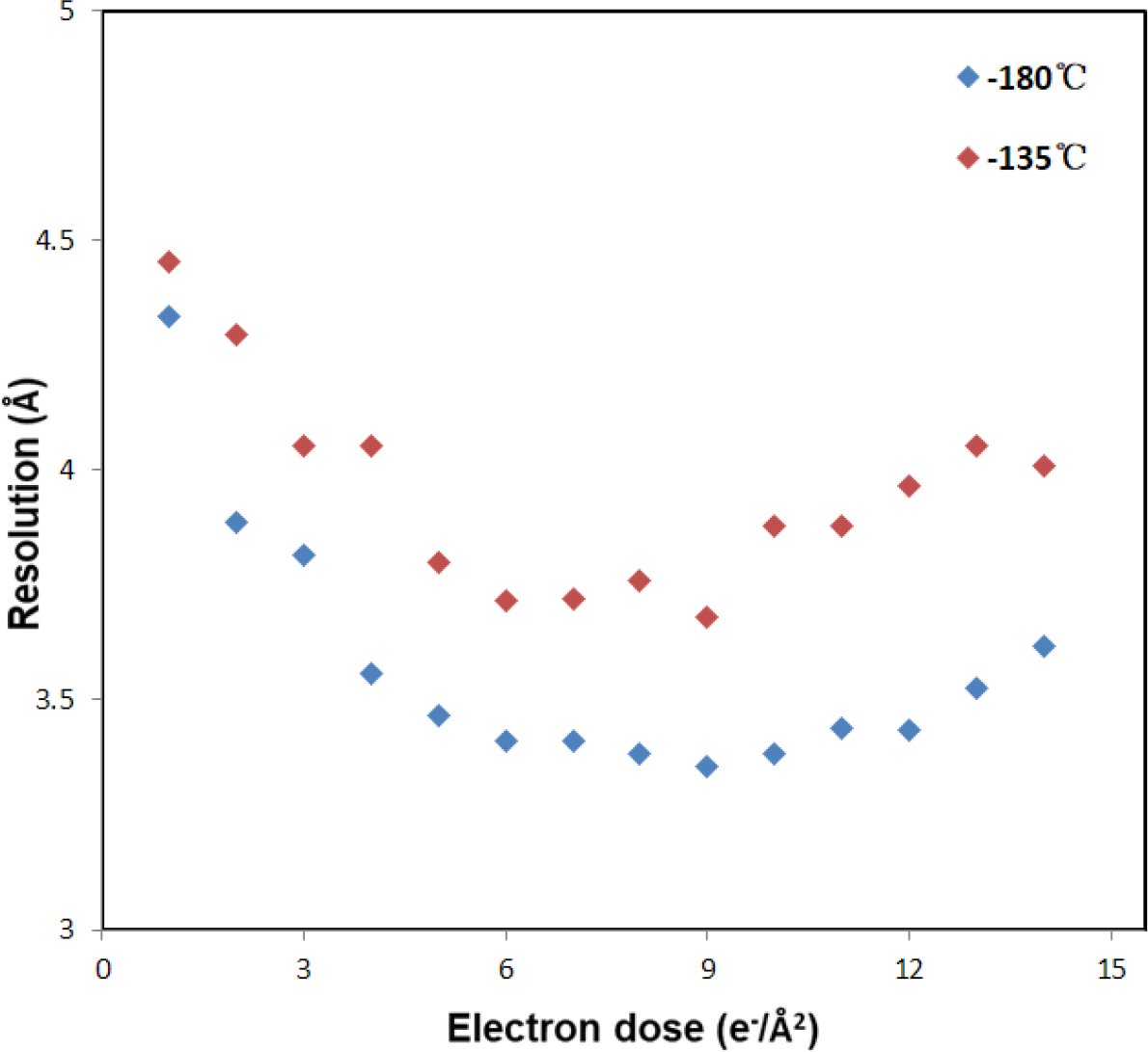
Resolution of the per-frame reconstructions of a virus with diameter of ~35 nm frozen at different temperatures. Each frame was exposed by 1 e^−^/Å^2^. The resolution of the per-frame reconstruction from the initial five frames decreased for the sample frozen at −180 °C (blue). Such a decrease became smaller for the sample frozen at −135 °C (red). The resolution of the reconstruction of the first frame is ~1.1 Å lower than that of the best reconstruction for the sample frozen at −180 °C while it is ~0.8 Å for the sample frozen at −135 °C.

**Fig. S5.**
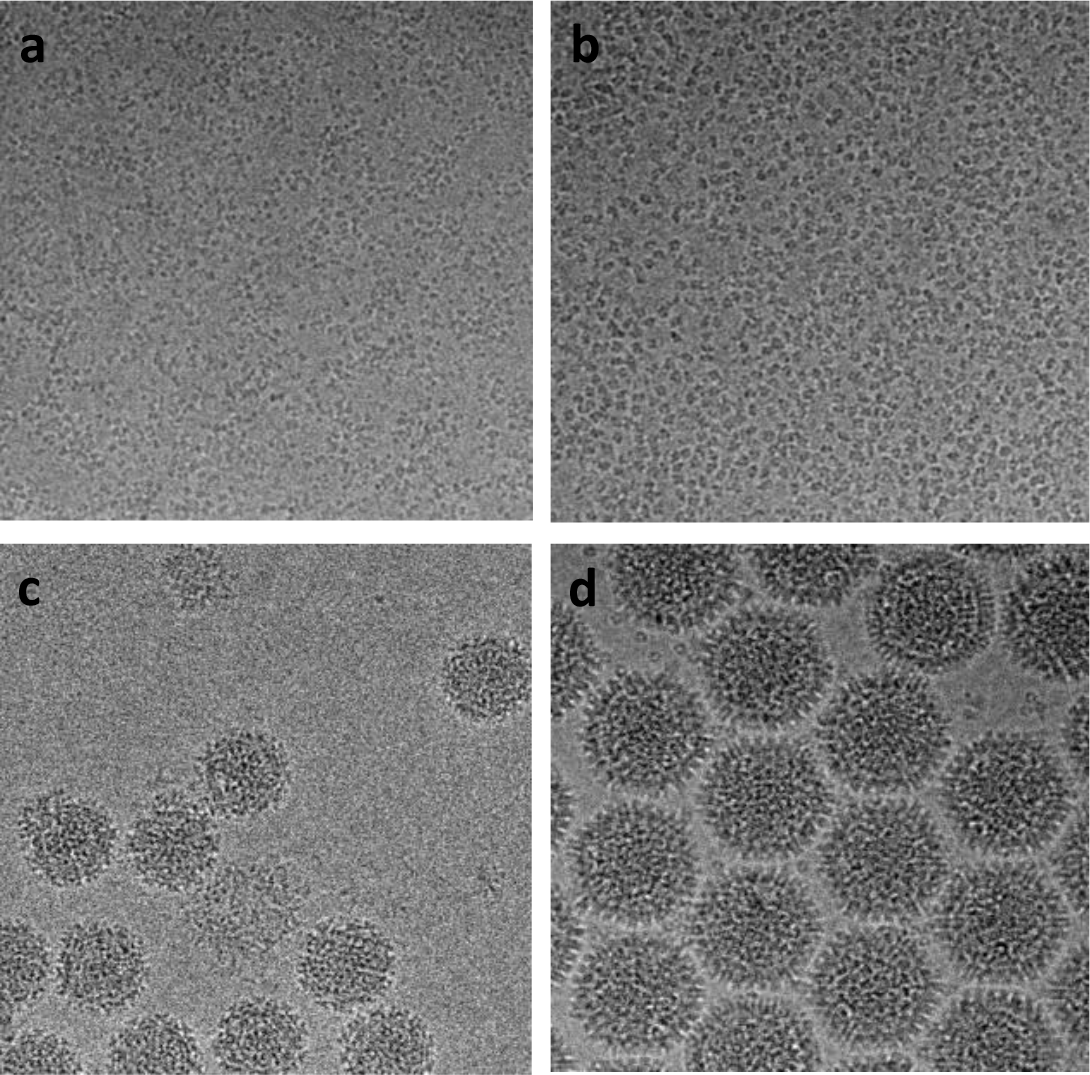
Samples vitrified at different temperatures. **a**, Hemoglobin (64 kDa) and **b**., Cas3-AcrF3 (a 130kD protein) frozen at −110 °C were embedded in the vitreous ice. **c**, A virus with diameter of ~55 nm and **d**, a virus with diameter of ~95 nm frozen at −130 °C were embedded in the vitreous ice.

